# Proteome modulation by opposite inotropic drugs in human engineered cardiac tissue revealed by topology-driven cross-modal integration

**DOI:** 10.64898/2026.08.25.746955

**Authors:** Doroteya K. Staykova, Danique Snippert, Hans J.C.T. Wessels, Robert Passier, Federica Conte

**Affiliations:** Multicore Dynamics Ltd, New Milton, United Kingdom; Applied Stem Cell Technologies, Department of Bioengineering Technologies, Cardiovascular Health Technology Centre, TechMed Centre, University of Twente, Enschede, the Netherlands; Department of Human Genetics, Radboud University Medical Center, 6525GA, Nijmegen, The Netherlands; Department of Anatomy and Embryology, Leiden University Medical Centre, 2311EZ, Leiden, The Netherlands

**Author notes:** Corresponding author: Dr. Federica Conte, Email address (University of Twente, NL), (Radboud University Medical Center, NL).

**Keywords:** (1) Cross-modal data integration, (2) Topological data analysis, (3) Drug-induced proteome modulation, (4) 3D engineered heart tissue, (5) Functional and molecular integrated drug screening

## Abstract

Engineered heart tissues (EHTs) represent an innovative platform enabling physiologically relevant *in vitro* evaluation of drug-induced cardiac responses. While functional characterization remains central to EHTs, molecular profiling is increasingly used to elucidate mechanisms underlying drug-induced phenotypes. Proteomics provides broad molecular characterization of drug responses at the protein level, yet the complexity, heterogeneity, and high dimensionality of proteomics datasets challenge conventional statistical approaches, which are not designed for cross-modal integration and streamlined multi-omics analysis. In this study, we developed an innovative framework based on topological data analysis (TDA) for the integration of large proteomics profiles and functional readouts to investigate system-level responses to drugs with opposing inotropic effects, epinephrine and doxorubicin. Samples were organized into a topological connectivity network according to multimodal similarity enabling simultaneous exploration of treatments, cardiac function and proteome alterations. Highly correlated features were then used for pathway enrichment analysis, which revealed strong similarities between the enrichment profiles associated with contractile force and epinephrine. These findings are consistent with the positive inotropic effect of epinephrine, whereas doxorubicin exhibited an opposing enrichment profile. Energy homeostasis, mitochondrial translation and proteostasis emerged as the major cellular processes displaying opposite associations with the two inotropic drugs, highlighting a link between cardiac contractility and perturbations in these processes. In conclusion, our TDA-based framework successfully integrated functional and proteomic data to uncover treatment-specific remodeling in EHTs, offering a modular and scalable approach that could be adapted to other *in vitro* organ models for systems-level mechanistic studies and next-generation drug development.

## INTRODUCTION

Organ-on-chip (OoC) models and engineered tissues have emerged as promising platforms for preclinical drug development because of their ability to recapitulate key aspects of human organ physiology *in vitro* while reducing reliance on animal models ^1–7^. In cardiac research, engineered heart tissues (EHTs) are three-dimensional, stem cell-derived cardiac models that enable the assessment of drug-induced perturbations in cardiac function, such as contractile force, making these models valuable for preclinical evaluation of drug efficacy and cardiotoxicity.

While functional characterization remains the cornerstone of cardiac tissue models such as EHTs, molecular profiling is becoming increasingly important for elucidating the mechanisms underlying functional phenotypes. Mass spectrometry (MS)-based proteomics is a rapidly expanding omics discipline broadly applied in therapeutic development and screening ^8^. Among the numerous types of proteomics, expression proteomics represents a powerful discovery tool that can help shed light on how drugs influence biological systems at the molecular level ^9–12^. This is particularly relevant in the development of cardioactive compounds, for which the therapeutic window is often narrow and off-target effects can have severe consequences ^13^. Cardiovascular physiology is regulated by a complex interplay of contractile proteins, signaling pathways, ion channels and metabolic regulators, all of which can be affected by pharmacological intervention. Recent advances in high-resolution MS-based proteomics have substantially enhanced our ability to characterize perturbations in the cardiac proteome ^14–16^. Beyond providing system-level insights into pharmacological effects and toxicity, expression proteomics can facilitate the identification of molecular targets and downstream pathways associated with these responses ^17–19^.

The integration of functional cardiac OoC models with high-resolution proteomics offers unprecedented opportunities to characterize drug responses across multiple biological layers. However, despite continued advances in tissue engineering and omics technologies to better replicate and quantify tissue complexity, analytical approaches have evolved more slowly. Conventional analytical methods largely rely on statistical approaches that were not designed to accommodate the complexity, heterogeneity, and high dimensionality of the datasets generated by technologies such as high-resolution expression proteomics ^20–24^. These challenges are amplified in OoC studies, where limited sample availability, small numbers of biological replicates due to biofabrication constraints, and biological heterogeneity complicate statistical inference ^25^. Moreover, these functional models allow the collection of different types of data, called *data modalities* (e.g. functional readouts, imaging data, and molecular profiles). Current data analysis pipelines do not allow direct cross-modal integration; rather, they are applied independently to each data modality, requiring additional analytical steps for the unified interpretation of heterogeneous datasets, such as functional and molecular readouts. To our knowledge, no reported approach has directly integrated high-dimensional omics data with phenotypic measurements without requiring prior modality-specific feature selection, feature balancing across modalities, or distributional harmonization. Such requirements can introduce information loss and preprocessing burden, hindering streamlined and reproducible cross-modal analysis ^26^.

These challenges motivate the adoption of alternative mathematical approaches capable of integrating heterogeneous data without requiring extensive modality-specific preprocessing or strong distributional assumptions. Topological data analysis (TDA) offers these advantages while also enabling the construction of a unified mathematical model that integrates and interactively visualizes multiple data modalities while preserving their underlying relationships ^27,28^. By leveraging concepts from algebraic topology, TDA focuses on the analysis of the “shape” of data, mapping how data points are organized and connected in space without relying on assumptions about linearity or distribution ^25,29^. In the context of cardioactive drug screening and development, TDA represents a powerful alternative approach to detect even subtle changes in protein expression patterns that may underlie key biological responses. Previous studies already demonstrated how TDA can successfully identify clusters of molecules (including proteins) that covary in response to treatment and highlight transitional states representing shifts from healthy to pathological conditions ^25,30^. This is particularly relevant to the interactions and interdependencies between proteins within tissues ^28^. Since pathophysiological changes often arise from complex, system-level disruptions in cardiac tissue, the ability to map these perturbations is critical in cardiovascular research, and preclinical biomedical research more broadly.

To capture system-level responses to cardioactive treatments with opposite functional effects in cardiac tissue, we employed a TDA-based framework designed to maximize information yield from heterogeneous datasets while remaining robust to small sample sizes. Building on our prior integrative work ^25^, we refined and expanded our analytical approach to mathematically integrate large-scale proteomic profiles with functional readouts from human EHTs. This framework allowed us to uncover the molecular processes directly associated with different functional perturbations induced by two drugs with opposite inotropic effects: epinephrine (positive) and doxorubicin (negative). Together, our findings demonstrate that the TDA-based framework integrates functional and proteomic data to uncover treatment-specific remodeling in EHTs, providing a scalable approach for cross-modal analysis of complex datasets in preclinical cardioactive drug development.

## MATERIALS & METHODS

### Cardiomyogenic differentiation

Cardiomyocytes (CMs) were differentiated from a commercial human induced pluripotent stem cell (hiPSC) line, GM25256 (Coriell Institute), following a previously established protocol (**Figure 1a**) ^25,31^. On day 13, metabolically active, spontaneously beating CMs were purified through glucose deprivation and lactate supplementation, using a glucose-free CM-specific medium supplemented with 5 mM sodium DL-lactate (60% w/w, Sigma-Aldrich), as previously described ^25^. From day 17 to day 20, CMs were cultured in CM medium supplemented with 4.5 mM glucose, prior to cryopreservation. Before freezing, the expression of cardiac troponin T was assessed in each batch by flow cytometry to evaluate batch purity.

**Figure 1.**
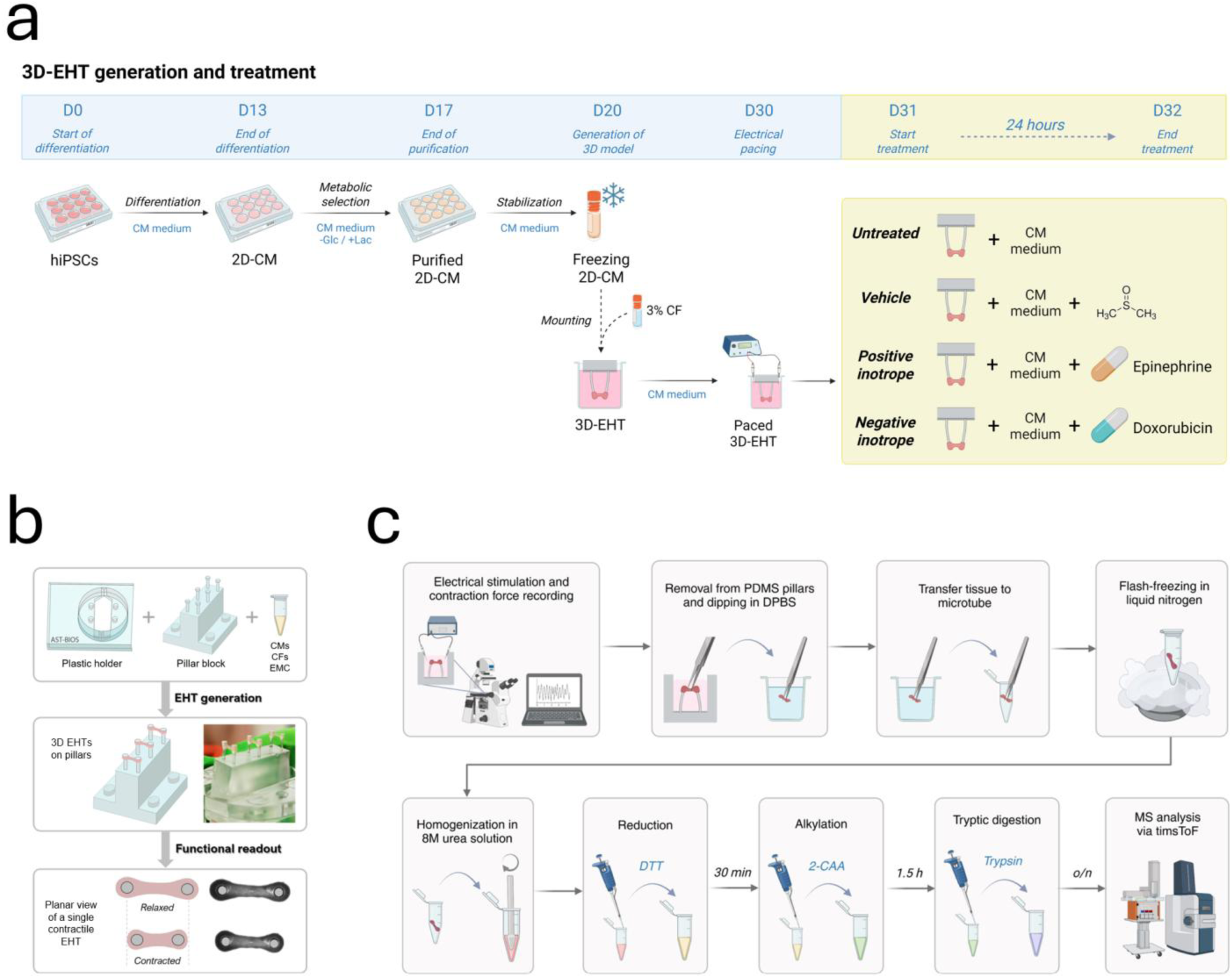
Functional EHT generation and sample processing protocol. (a) Timeline depicting the cardiomyogenic differentiation, followed by EHT generation and treatment. (b) Platform components and schematic representation of functional EHTs cast on flexible plastic pillars. (c) Sample processing protocol for EHTs, performed directly after video recording for FoC quantification. Abbreviations: 2-CAA, 2-chloroacetamide; DTT, dithiothreitol; PDMS, polydimethylsiloxane; DPBS, Dulbecco’s phosphate-buffered saline (without calcium and magnesium).

### Generation of functional 3D engineered heart tissues

For the generation of 3D-EHTs, commercially available human adult cardiac fibroblasts (CFs) (C-12375, Promocell) were acquired and used for tissue generation, as described previously ^25,31^. In brief, CMs and CFs were thawed on ice, counted and combined at a final composition of 100:3 CMs:CFs (**Figure 1a,b**). The cell suspension was then mixed with an extracellular matrix mixture of defined composition ^25^, containing fibrinogen and thrombin (100U/mL, Sigma), the latter added immediately before casting the mixture into gelatin molds to initiate gel polymerization around plastic pillars ^32^. The platform used has been previously described ^31,32^, and it contains three sets of pillars for each holder, to enable the culture of three replicates in the same well of a 12-well plate (**Figure 1b**). The 3D-EHTs were maintained at 37 °C and 5% CO₂ in a humidified incubator for 24 hours before refreshing with CM culture medium. Subsequently, the CM culture medium was refreshed every 48 hours until day 9 post-tissue casting (**Figure 1a**).

### Treatment with inotropic drugs

On day 9 after tissue formation (corresponding to day 29 in the general timeline in **Figure 1a**), the EHTs were divided into five groups. Group 1 included untreated tissues (*n=3*) harvested and flash-frozen in liquid nitrogen immediately after pacing and video recording for force of contraction (FoC) measurement, representing the timepoint 0 of the experiment (baseline). The remaining 12 tissues (*n=3* for each of the four conditions) were moved into the wells of a new plate, each containing 2 mL of CM medium supplemented with: nothing (untreated tissues, group 2), 7.05 μM DMSO (1:2,000 dilution, vehicle control condition, group 3), 1 μM epinephrine (EPI, 1:10,000 dilution, group 4) or 5 μM doxorubicin (DOXO, 1:2,000 dilution, group 5). After 24 hours of treatment exposure, the 12 tissues were paced and recorded for FoC measurement, then immediately removed from the pillars and flash-frozen in liquid nitrogen for subsequent protein extraction.

### Image-based contractile force measurement

To evaluate the cardiac function (i.e. FoC) of the treated 3D-EHTs, real-time video recordings were acquired during electrical pacing. Tissues were paced at 2 Hz using a custom-designed electrical stimulator connected to platinum electrodes (Advent Research Materials), delivering biphasic pulses (10 ms duration, 4-5 V/cm) for a total of 20 seconds, as previously reported ^33^. Recordings were captured using a Nikon ECLIPSE Ti2-E inverted microscope coupled with a Prime BSI high-speed camera (Photometrics), operating at 100 frames per second and 2X magnification. All measurements were conducted under physiological conditions (37°C, 5% CO₂) to preserve tissue functionality.

Contractile force was computed from video data using the EHT Analysis software ^33^, which translates pillar deflections into quantitative force readouts. Functional readouts were collected at two stages: before treatment (baseline, 0h, *n=3*) and after 24 hours of exposure to drugs or control condition (*n=3* per condition, 4 conditions in total). Besides using the 24h force values, the baseline FoC measurements before treatment (0h) were also collected for all EHTs to calculate the relative force change over time.

### Protein extraction from individual EHTs

Immediately after video recording for FoC quantification 24 hours after treatment initiation (**Figure 1c**), each tissue was removed from the pillars using tweezers, transferred to labeled microtubes and flash-frozen in liquid nitrogen and stored at-80°C till extraction. On the day of extraction, the frozen tissues were ground using single-use plastic pestles for 1.5 mL microtubes (Cyvta) in 25 uL of 8 M Urea/10 mM Tris pH 8.0 until complete tissue homogenization (approximately 40 sec/tissue). Then, 25 μL reduction buffer, consisting of 10 mM dithiothreitol (Sigma) in ultrapure water (HPLC-super gradient grade, VWR Chemicals), was added to each sample and incubated at room temperature for 30 minutes. Next, 25 µL of alkylation buffer, consisting of 50 mM 2-chloroacetamide (Sigma) in 50 mM ammonium bicarbonate (Sigma), was added to each sample and incubated for 1.5 hours at room temperature in the dark. Lastly, 50 μL of 50mM ammonium bicarbonate were added to each sample and mixed by vortex for 5 seconds, followed by addition of 3 μL of trypsin (0.4 µg/µL protein, Promega). The samples were then incubated for 15 hours at 37°C. The following day, after spectrophotometric quantification of peptide concentration, the samples were loaded onto EVOtips (Evosep), according to manufacturer’s instructions, prior to MS analysis.

### Proteomics analysis and data acquisition

Tryptic digests from EHT homogenates were analyzed by nanoflow liquid chromatography (Evosep One, Evosep Biosystems) coupled online to a trapped ion mobility spectrometry quadrupole time-of-flight mass spectrometer (timsTOF Pro2, Bruker Daltonics) via a CaptiveSprayer nanoflow electrospray ionization source (Bruker Daltonics). Tryptic peptides were separated by C18 reversed phase liquid chromatography (Evosep EV1109 30SPD performance column; 80 mm length x 0.150 mm internal diameter, 1.5 µm C18AQ particles) using the pre-programmed 30 samples per day (30SPD) Evosep One method. The MS was operated in positive ionization mode using the default long gradient data-independent acquisition / Parallel Accumulation SErial Fragmentation (dia-PASEF) instrument method: 0.6-1.6 1/K0 mobility range, 100-1700 *m/z* mass range, 100 ms accumulation time, 100 ms ramp time, 26 Da mass width, 1 Da mass overlap, 32 mass steps per cycle, 0 mobility overlap, 1 mobility window.

Acquired spectra were streamed directly to ProteoScape (v.2024, Bruker Daltonics) for protein identification and label-free quantitation against a *H. sapiens* protein sequence database (Uniprot, reviewed sequences including isoforms and canonical sequences downloaded in August 2024), using the following settings: Spectronaut v19 directDIA+ (Fast) workflow, 0.2 precursor PEP cutoff, 0.01 precursor Q-value cutoff, 0.01 protein Q-value cutoff global, 0.01 protein Q-value cutoff, 0.75 protein PEP cutoff, full tryptic specificity, up to 2 missed cleavages allowed, carbamidomethyl (C) as fixed modification and oxidation (M) as variable modifications. Protein group-specific peptides were used for quantitation. All details regarding the settings used for the proteomics data annotation are reported in Supp. File 1.

### Proteomics data processing

After PaSER data preprocessing and extraction, analytical replicates (triplicate injections) were averaged, based on the high reproducibility observed between these replicates and the low variability in the mass spectrometry measurements. The original scale of protein abundance values was retained (no log2-transformation performed) to preserve the intrinsic geometry of the proteomics space prior to TDA.

For downstream analysis, protein abundances were organized into an annotated data matrix (proteins as columns and samples as rows), with sample metadata arranged as observation-level annotations within an *AnnData* object ^34^. Data processing steps were implemented using custom-built code developed in Python v3.10.

### Topological graph modelling across molecular and functional scales in EHTs

We first translated the sample-centric experimental design into a mathematical space, where each protein, functional, and treatment variable defined a dimension, and its measured value for each sample defined the corresponding coordinate. The EHT samples therefore formed a point cloud sampled from this multimodal space. We then used our previously described TDA-based framework to capture the topological structure of the sampled space and represent its underlying “shape” as a graph. Molecular and functional measurements were used to define topological lenses that map the multimodal space within the *Mapper* algorithm ^35^. Lens values were used to partition the sample space into overlapping intervals, within which samples were clustered into network nodes. Nodes were connected when they shared one or more samples, capturing the local structure and connectivity of the data space ^27,36,37^. The resulting graph is a unified topological model of the EHT system, herein referred to as topological connectivity network (TCN).

A key step was defining which data layers to include in the TCN and which ones to retain as metadata for subsequent interpretation. To assess proteomic variation and sample relationships, we evaluated four unbiased low-dimensional projections generated with principal component analysis (PCA), principal coordinates analysis (PCoA) ^38^, uniform manifold approximation and projection (UMAP) ^39^, and potential of heat-diffusion for affinity-based trajectory embedding (PHATE) ^40^. PCoA was selected for subsequent analysis because it preserves pairwise sample relationships in a low-dimensional space while allowing flexible selection of distance metric ^38^. Only human proteins with complete annotations were included in the downstream analysis. PCoA was applied to a pairwise correlation distance matrix computed from protein abundance values using the *scikit-bio* Python library (v.0.7.1.post1) ^41^.

The tissue contractile force (i.e. FoC) and the first two principal coordinates derived from proteomics PCoA were used as topological lenses, whereas treatment conditions served as metadata. This design enabled integration of heterogeneous data modalities while allowing treatment-associated proteome and functional responses to be explored in a data-driven, holistic manner. The resulting TCN was constructed using the *giotto-tda* package ^42^.

### Node-based correlation analysis for inferring in vitro treatment-induced proteomic changes

The interactive, unified mathematical model, the TCN, which offers both intuitive visualization and direct access to the underlying diverse and high-dimensional data, represented the basis for downstream analyses. To translate the qualitative information obtained by the TCN exploration into quantitative data that can be further analyzed, for instance via standard statistics, we leveraged an approach recently benchmarked, namely topological node-level enrichment (TNE) ^25^. In brief, for each TCN node containing clustered samples, we calculated node values based on the data type as follows. (1) For proteins, we computed the mean values of corresponding protein abundances (see *Proteomics data processing*). (2) For FoC, measurements were averaged per node. (3) For categorical metadata, such as treatment type, the node value represented the percentage of samples within the cluster exhibiting the associated category. Subsequently, a node-level dataset consisting of functional and molecular data, along with experimental metadata across the entire topological network was constructed. To assess the strength and direction of associations across data modalities, Pearson correlation was calculated using the node values. Proteins were retained for downstream pathway enrichment analysis only if they showed statistically significant correlations (*p* < 0.05) with absolute correlation coefficients > 0.7 to restrict pathway-level interpretation to strong and statistically robust associations.

### Pathway enrichment analysis, selection and visualization

Pathway enrichment analysis was performed using the Reactome Pathway Database (reactome.org, August 2026). Separated Uniprot accession lists of negatively-and positively-correlating proteins were loaded in the Reactome Analysis Tools interface, setting the ‘project to human’ option and disabling the ‘include interactors’ option (to restrict enrichment to manually curated pathway members).

Next, the resulting pathways were sorted and filtered based on the uncorrected p-value (*p* < 0.05) and estimated false discovery rate FDR (*FDR* < 0.1) for visualization purposes. The resulting significantly associated pathways were visualized as weighted Voronoi treemaps using FoamTree (Carrot Search, FoamTree, Version 3.5.7, 2025; carrotsearch.com/foamtree/). Pathway p-values were transformed as − log_10_(*p*) and used to determine the size of Voronoi graph tiles. FDR values were transformed as − log_10_(*FDR*) and rescaled to [0,1] using min-max normalization with a fixed lower bound of 0 and an upper bound defined as max(−log_10_(*FDR*), −log_10_(0.05)), ensuring that the significance threshold (*FDR* = 0.05) was represented in the color scale. The normalized values were used for continuous color mapping. Yellow and red color gradients were selected to represent positive and negative correlations, respectively. A custom workflow implemented in Python 3.10 was used to process.csv pathway output files and generate interactive.html visualizations of the Voronoi treemaps based on FoamTree.

To identify shared pathways across treatments, pathway sets associated with FoC, EPI, and DOXO were matched based on the pathway identifier. Shared pathways were visualized using chord diagrams. Ribbon thickness was calculated as − log_10_(*FDR*), with thicker ribbons corresponding to lower FDR values. Ribbon color indicated the direction of association (yellow, positive; red, negative). Chord diagrams were generated using the ‘chordDiagram’ function from the R package *circlize* (v0.4.15).

## RESULTS

### 3D-EHTs showed opposite functional trends after treatment with positive and negative inotropic drugs

The set of 3D EHTs was successfully generated from a healthy, commercially available hiPSC line, and cultured for 9 days prior to electrical pacing and treatment (**Figure 1a,b**). On day 9, the EHTs were paced, and videos were recorded to quantify the baseline FoC in all the tissues. The EHTs were then divided into five groups. One group (3 EHTs/well) was collected immediately as the untreated, 0h group. The remaining tissues were divided into four groups and left untreated or treated with either DMSO (7.05 μM), EPI (1 μM) or DOXO (5 μM). All treated EHTs survived the 24h incubation with the treatments and showed contraction, although the latter was differentially affected by the treatment, as shown in **Figure 2a**. Video analysis for FoC quantification (Supp. Dataset 1) showed comparable force levels in untreated and DMSO-treated EHTs, with both displaying slightly lower force than untreated tissues measured at 0h. Compared to untreated and DMSO-treated EHTs, EPI-treated EHTs displayed increased FoC, indicating a positive inotropic effect, whereas DOXO-treated EHTs showed a clear decrease in FoC, indicating a negative inotropic effect (**Figure 2b**). The 24-hour time window was selected to allow detectable variation in FoC without compromising tissue integrity or viability of the tissues. After recording, each EHT was directly collected for processing, tryptic digestion and MS-based proteomics analysis. Overall, EHTs across all conditions showed highly comparable protein-group detection (**Figure 2c**). Following label-free quantification (LFQ) with match-between-run (MBR) analysis, 7,124 protein groups were detected across the EHT sample set (Supp. Dataset 2).

**Figure 2.**
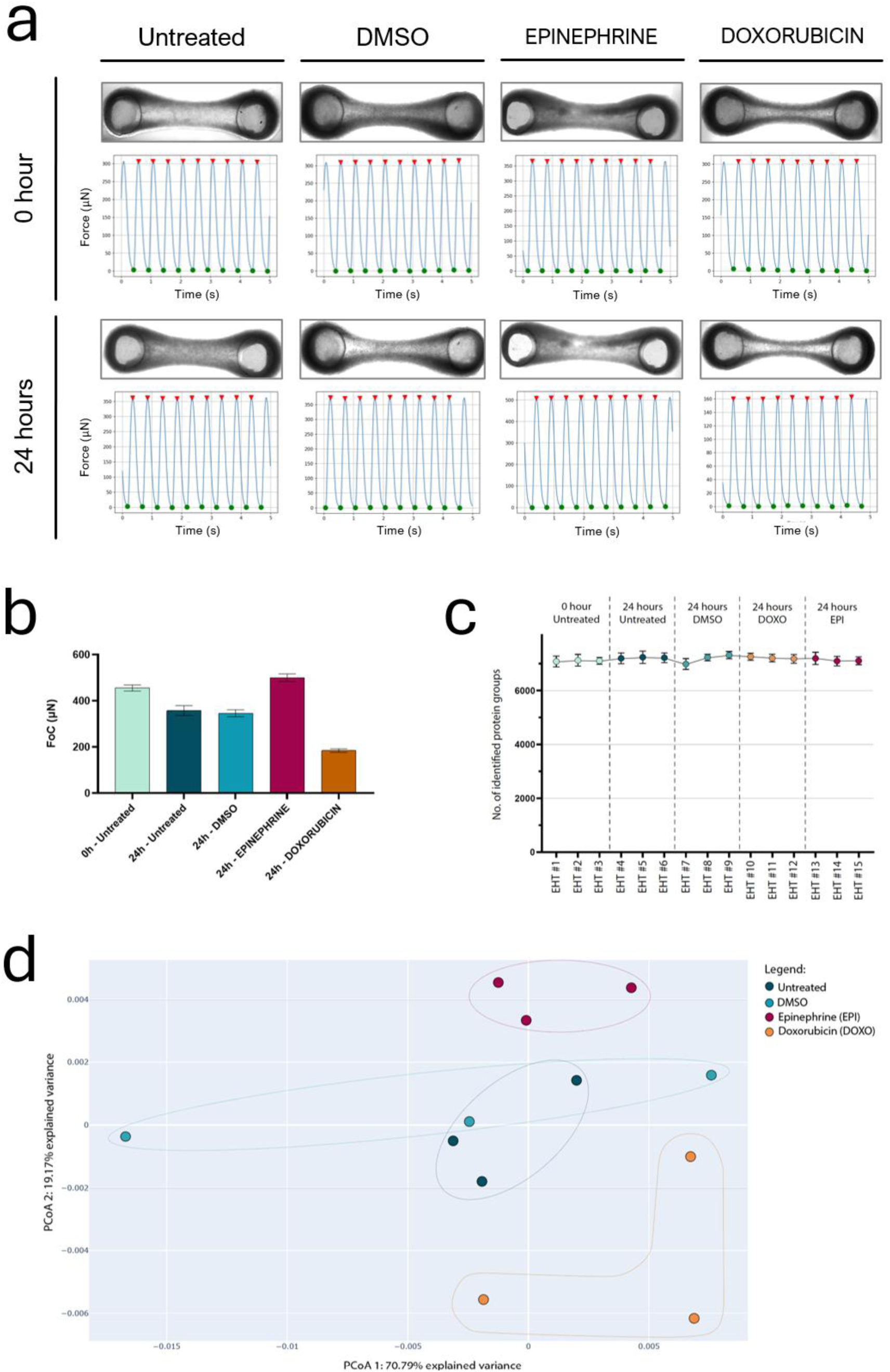
Overview of functional readout and proteomics results. (a) Representative force curves graphs obtained from untreated and treated EHTs. Above each graph, optical microscopy images show EHT morphology before and after treatment. One tissue is shown as an example, although three tissues were measured per condition. (b) Bar graph showing the FoC calculated in each set of EHT triplicates (0h vs 24h, treated vs untreated, n=3). The bar indicates the standard deviation. (c) Graph representing the total number of protein groups (PG) detected for each EHT before LFQ-based MBR. (d) Principal coordinate analysis (PCoA) of the proteomics profiles resulting from the EHTs collected at 24 hours from treatment initiation. Each dot represents the average of the protein levels from the three replicate EHTs (*n=3*).

### Topological data analysis of proteomic data structure revealed time-and condition-dependent effects

After data preprocessing, four unbiased dimensionality-reduction methods were applied to evaluate patterns of proteome variation and their effects on sample relationships: PCA, PCoA ^38^, UMAP ^39^ and PHATE ^40^. The four projections were applied to and evaluated using protein abundance values without log-transformation. Unlike conventional statistical workflows, TDA seeks to reveal the connectivity structure and topology of the data space. Nonlinear transformations, such as log_2_ scaling, can alter pairwise distances between samples, thereby affecting sample relationships and connectivity patterns, consequently altering the shape of the data captured by TDA.

All four approaches revealed a strong time-dependent effect, with the 0-hour baseline samples forming a distinct cluster that was clearly separated from the 24-hour samples (**Supp. Figure 1**). This pronounced shift in proteomic state indicated that time of collection was a major source of variation, exceeding the variation observed across treatments. Because this strong time-associated change in state could confound the interpretation of treatment-induced proteomic changes, 0h baseline samples were excluded from treatment-focused analyses.

To further characterize sample relationships and assess treatment-associated variation, we performed PCoA using a correlation-distance metric, thereby preserving pairwise similarities between samples in a low-dimensional embedding (**Figure 2d**). The first two principal coordinate axes accounted for 70.79% and 19.17% of the total variance, respectively. The projection revealed a clear separation of samples according to treatment. The three EPI-treated replicates formed a compact cluster characterized by high values along PCoA axis 2, whereas the DOXO-treated samples clustered at high values along PCoA axis 1 and low values along PCoA axis 2. Untreated and DMSO-treated samples occupied an intermediate region of the embedding and partially overlapped, consistent with the expected similarity between the untreated and vehicle-control (DMSO) conditions.

The separation of the four treatment groups indicates that treatment-specific proteomic changes are the primary drivers of the variation captured by the PCoA embedding. The first two PCoA coordinates were therefore selected as filter functions (topological lenses) for downstream TCN construction.

### Multi-layer TCN revealed distinct regimes in the functional-molecular landscape of EHTs

Using the PCoA coordinates as filter functions, we constructed a multilayer topological connectivity network (TCN) integrating contractile force measurements, large-scale proteomics, and treatment categories from EHTs (**Figure 3a**). Table 1 summarizes the information layers incorporated into the mathematical model. The first two topological lenses provided a molecular view based on the first two PCoA coordinates, capturing the major axes of proteomic variation (**Figure 2d**). FoC data were included as a third topological lens to represent tissue function, with reduced force observed in DOXO-treated samples compared with the other conditions (**Figure 2b**).

**Figure 3.**
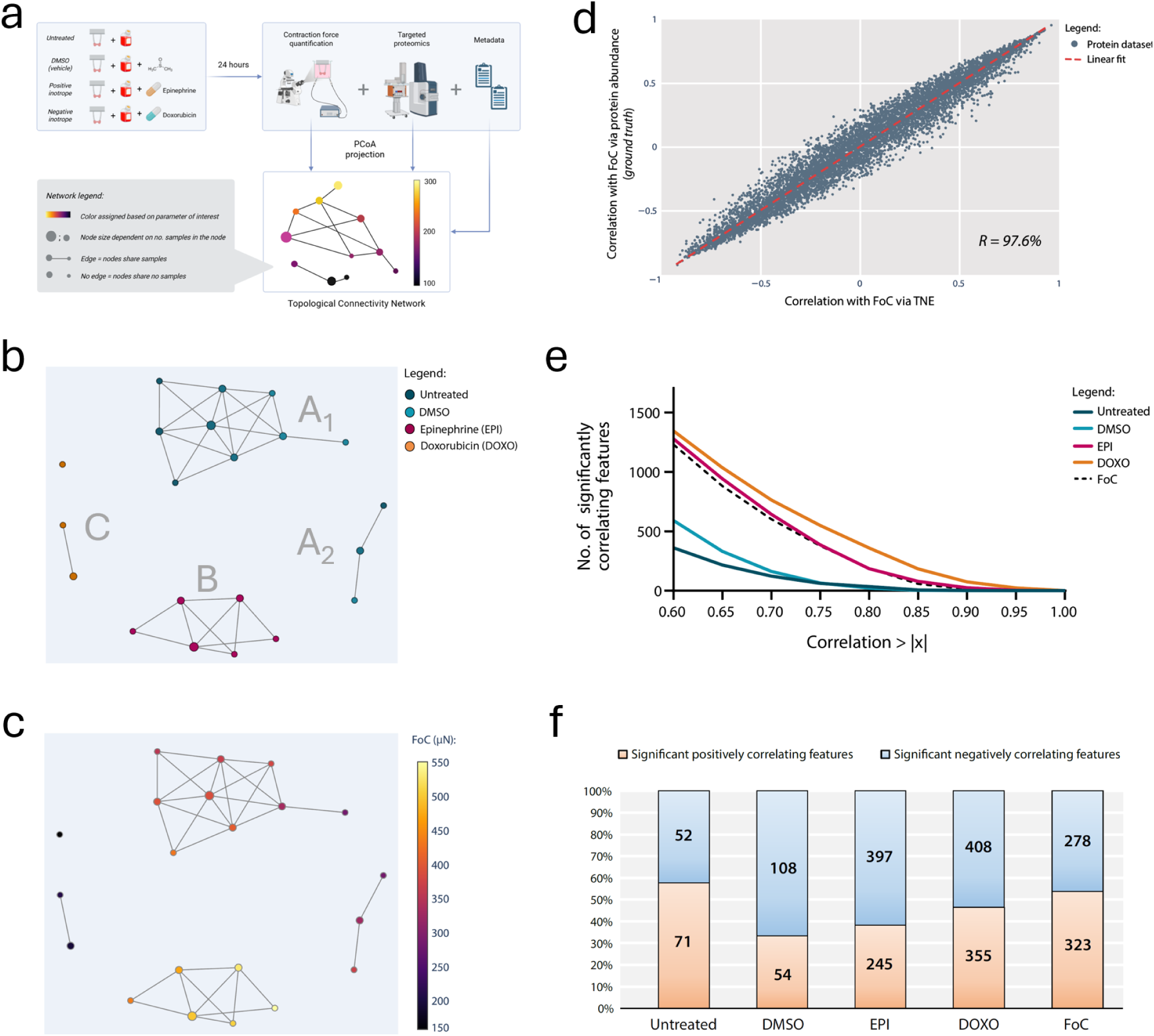
Topological connectivity network (TCN) as the unified mathematical model integrating functional readout, proteomics profiling and metadata. (a) Schematic representation of the integration framework and TCN legend. (b) TCN colored based on condition. Each node includes different samples (see Methods), with treatment conditions assigned to distinct colors. Node colors reflected the relative proportion of samples from each condition, with colors blended accordingly for mixed nodes. Five separate elements appear in the TCN. A large upper cluster (A_1_) encompasses untreated and DMSO-treated samples, flanked by a 3-node cluster on the right (A_2_) also containing untreated and DMSO-treated samples. Below, a 6-node cluster contains all EPI-treated samples (B). Lastly, on the left a 2-node cluster (C) and an isolated node contain the DOXO-treated samples. (c) TCN colored based on FoC, showing that the nodes in the lower cluster (B) containing EPI-treated samples show the highest FoC values, while the 2-node cluster (C) and single node containing DOXO-treated samples display the lowest FoC values. The large upper cluster (A_1_) and lateral 3-node cluster (A_2_) containing the untreated and DMSO-treated samples display intermediate FoC values. The interactive version of the TCN, in which the user can change the node coloring based on different parameters (specific samples, metadata, etc.) is available as Supp. File 2. (d) Proteome-wide benchmarking of our approach demonstrates robust agreement between TCN-based inference and abundance-based correlation for estimating strength of associations with FoC (R=97.6%). (e) Line graph summarizing the total number of significantly correlating features for each condition at different correlation thresholds (x): |correlation| > 0.6, 0.65, 0.7, 0.75, 0.8, 0.85, 0.9, 0.95, 1. (f) Bar graph summarizing the significantly (p < 0.05) positively and negatively correlating features for each condition.

**Table 1.** Summary of information layers and their encoding in the *Mapper*-derived TCN model.

| Information included as *topological lenses* | Information included as *metadata* |
| --- | --- |
| Two-component PCoA projection of 7,124 protein profiles | Treatment group |
| Contractile force of 3D-EHTs | Biological replicate (within treatment group) |
|  | Relative change in contractile force |

In brief, the TCN is a network graph in which nodes represent clusters of samples with similar profiles. Samples within each lens’ interval are grouped into network nodes, which are connected by an edge when they share one or more samples, thereby capturing the local structure and connectivity of the dataspace (**Figure 3a**).

Visual exploration of the TCN identified 4 connected components, with samples clearly segregating according to treatment condition (**Figure 3b**, Supp. File 2). The largest 9-node component (A_1_), together with the adjacent 3-node component (A_2_), comprised all untreated and DMSO-treated samples. In the lower region of the TCN, a 6-node component (B) consisted exclusively of EPI-treated samples. Finally, on the left side of the network, a small 2-node component (C) and an isolated node contained only DOXO-treated samples. Coloring the TCN according to FoC further reinforced this organization (Figure 3c). The lowest force values were observed in the DOXO-associated region (component C and the isolated node), indicating a network area associated with a pronounced functional shift, consistent with the negative inotropic effect of DOXO. In contrast, the highest force values were concentrated within component B, which contained the EPI-treated samples, consistent with the positive inotropic effect of EPI. Intermediate force values were observed across components A_1_ and A_2_, which contained untreated and DMSO-treated samples, consistent with the preserved contractile function expected under these control conditions.

Overall, the organization of the TCN successfully and simultaneously reflected treatment conditions, contractile force, and proteomic variation, thus providing a systems-level representation of treatment-induced responses in EHTs.

### Node-level correlation analysis enables stratification of inotropic drug effects across functional and proteomics layers

Building on our previous work ^25^ and the structure of the TCN described above, we investigated how EPI, a positive inotrope, and DOXO, a negative inotrope, shaped coordinated molecular and functional responses in EHTs. Because each TCN node represents a cluster of samples with similar profiles, the network partitions the multimodal data into locally coherent subsets. For each node, protein abundances and FoC were derived as node-level descriptors (**Supp. File 3, Supp Dataset 3**). Cross-modal associations between protein abundance and FoC were then quantified across nodes using Pearson’s correlation coefficient. Node-level associations between protein abundance and FoC showed strong agreement with correlations derived from the raw data across all detected protein groups (**Figure 3d**), supporting the robustness of the TCN framework for inferring cross-modal relationships.

Next, we quantified treatment composition of each node as the proportion of samples exposed to each condition (see Methods). Drug-specific associations with functional output and protein abundance were subsequently assessed across nodes using Pearson correlation (**Figure 3e,f** and **Supp. Dataset 4**). Only proteins exhibiting strong associations (|correlation| > 0.7) were selected for downstream pathway analysis (Supp. Table 1).

An additional advantage of our topological node-level enrichment (TNE) framework is its ability to map variables from different data modalities onto a common graph representation. Both numerical and categorical metadata can therefore be quantified at the node level, enabling their direct comparison using TNE-based correlations or other mathematical operations. For example, heterogeneous variables such as relative FoC changes and treatment conditions were mapped onto the nodes (**Supp. Figures 2 and 3**), enabling their relationships to be quantified.

These findings demonstrate that our approach is effective in unifying heterogeneous data modalities and provides a biologically interpretable map for investigating treatment responses in EHTs.

### Pathway enrichment of TDA-derived features reveals opposite pathway perturbations associated with epinephrine and doxorubicin

After applying a correlation threshold (|correlation| > 0.7), significant features were separated into positively and negatively correlated groups, and pathway enrichment analysis was performed using the Reactome platform (results are reported in Supp. File 4). Although informative for facilitating the interpretation of hundreds of significant features, pathway enrichment analysis has limitations that should be considered. Reactome pathways are hierarchically organized and often share annotated proteins, meaning that multiple significant pathways may be driven by the same set of proteins and consequently overlap in their biological annotations.

We first examined the pathway enrichment profiles associated with FoC, used here as the functional ground truth for contractile performance (**Figure 4a, d**). Overall, the pathways associated with FoC showed a striking similarity to those associated with EPI treatment (**Figure 4b,e**), consistent with the positive inotropic effect of EPI and its concordant relationship with the contractile force. In contrast, DOXO displayed a nearly opposite enrichment pattern (**Figure 4c,f**), with several pathways, particularly those linked to energy metabolism, showing a reversal in their direction of association relative to FoC and EPI.

**Figure 4.**
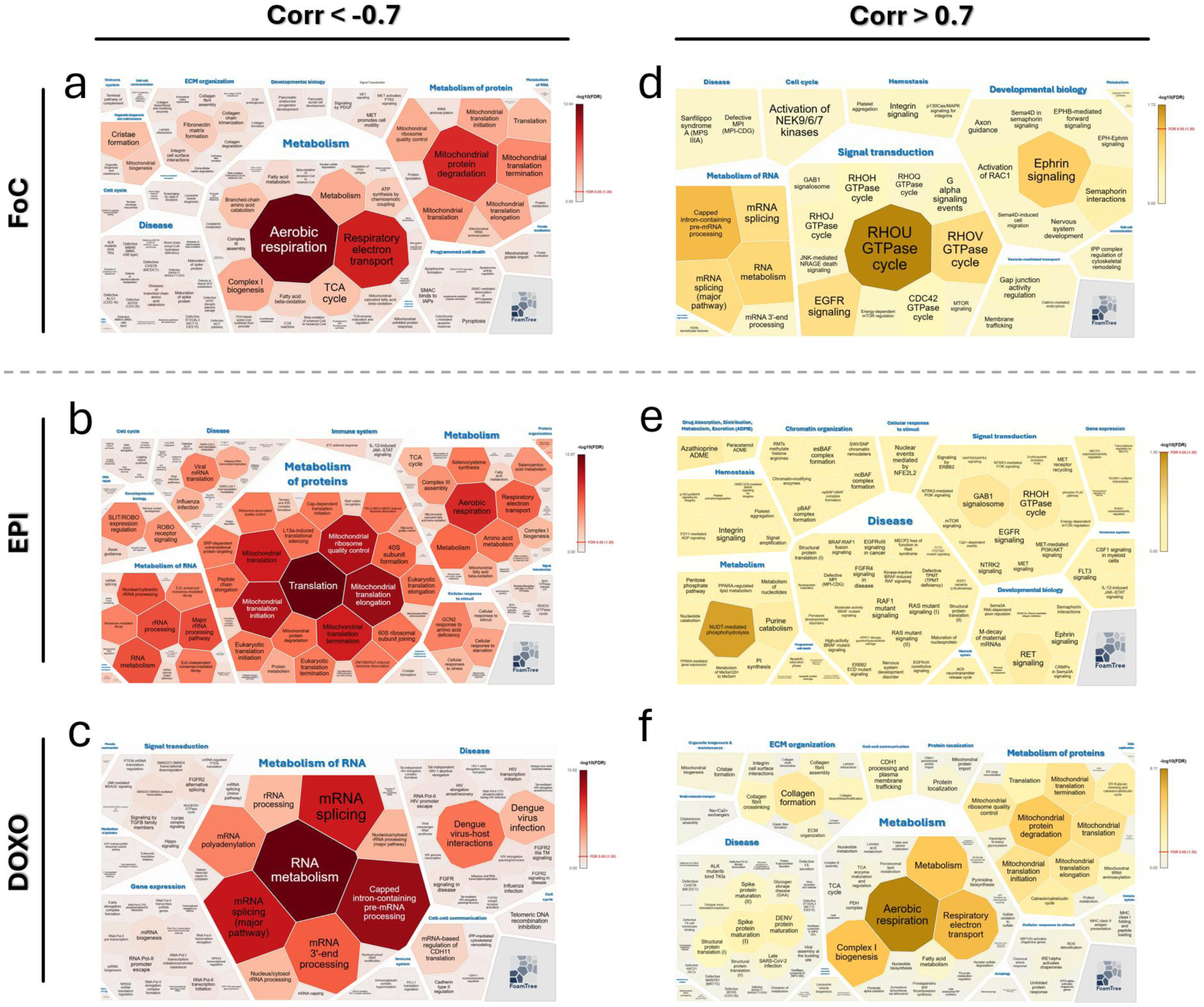
Customized weighted Voronoi diagrams summarizing Reactome pathway enrichment analysis. Weighted Voronoi diagrams visualize significantly enriched Reactome pathways identified from proteins showing strong correlations with FoC (ground truth) and the two drug-based treatment conditions. Red graphs depict pathways enriched among features exhibiting significant negative correlations (Pearson correlation < −0.7, p < 0.05) for FoC (a), epinephrine (EPI; b), and doxorubicin (DOXO; c). Yellow graphs depict pathways enriched among features exhibiting significant positive correlations (Pearson correlation > 0.7, p < 0.05) for FoC (d), EPI (e), and DOXO (f). Polygon size is inversely proportional to the raw enrichment *p*-value, while the color gradient encodes the FDR, with darker colors corresponding to lower FDR values. Pathway enrichment results for the untreated and DMSO conditions are presented in Supp. Figures 4 and 5. To improve visualization, high-resolution, zoomable PDF versions of all Voronoi graphs are provided as Supp. Files 5-14. The results of the enrichment analysis for all conditions (including the R-HSA indexes of the enriched pathways) are reported in Supp. File 4.

Specifically, negatively correlating features with FoC were predominantly associated with energy metabolism and mitochondrial processes. The most prominent enriched pathways included aerobic respiration, respiratory electron transport, fatty acid (FA) metabolism, tricarboxylic acid (TCA) cycle, mitochondrial protein degradation, and mitochondrial translation, indicating that proteins associated with mitochondrial bioenergetics and proteostasis were strongly associated with contractile function (**Figure 4a**). Positively correlated features were instead enriched in signal-transduction pathways, particularly RHO and RHOV GTPase cycles, EGFR signaling, RAC1 activation, and related intracellular signaling processes (**Figure 4d**). These findings highlight the link between contractile function, the regulation of intracellular signaling and the adaptation of (mitochondrial) energy metabolism. These findings suggest a potential, direct link between the contractile function and the coordinated regulation of mitochondrial metabolism and intracellular signaling.

The enrichment profile associated with EPI closely mirrored that observed for FoC. Negatively correlated features were predominantly associated with protein metabolic processes, including mitochondrial translation and mitochondrial ribosome-associated pathways, and energy metabolism, including aerobic respiration, respiratory electron transport, FA β-oxidation and TCA cycle (Figure 4b). Additional enrichment of pathways related to RNA metabolism was observed, underlying a broader remodeling of cellular proteostasis. Positively correlating EPI features were enriched predominantly in nucleotide metabolism and cellular signal transduction, including RHO and RHOV GTPase cycles, EGFR signaling, RAC1 activation, and other GTPase-mediated processes (**Figure 4e**).

Lastly, DOXO exhibited a nearly opposite enrichment profile compared to FoC and EPI. Negatively correlating features following DOXO treatment revealed prominent enrichment of RNA metabolism and post-transcriptional regulation, including mRNA splicing, mRNA 3′-end processing and pre-mRNA processing, together with pathways related to viral infection (Figure 4c). Conversely, positively correlating features were strongly enriched in mitochondrial and metabolic pathways linked to energy homeostasis, including aerobic respiration, respiratory electron transport and FA β-oxidation. Pathways linked to mitochondrial translation and mitochondrial protein homeostasis also resulted strongly positively associated with DOXO. Additionally, enrichment of pathways linked to extracellular matrix organization and collagen-related processes suggested alterations occurring at a structural level. Comparison of the enrichment profiles across FoC and the treatment conditions further highlighted a subset of pathways exhibiting an opposite direction of association between the positive inotropic FoC/EPI conditions and DOXO (**Figure 5**). The strongest overlap was observed among pathways negatively associated with both FoC and EPI and positively associated with DOXO, where energy homeostasis, mitochondrial translation and mitochondrial protein homeostasis represent the most prominent shared functional categories (**Figure 5a**). This pattern suggests that these interconnected metabolic and mitochondrial processes are closely associated with changes in cardiac contractile function induced by treatments with opposing inotropic effects.

**Figure 5.**
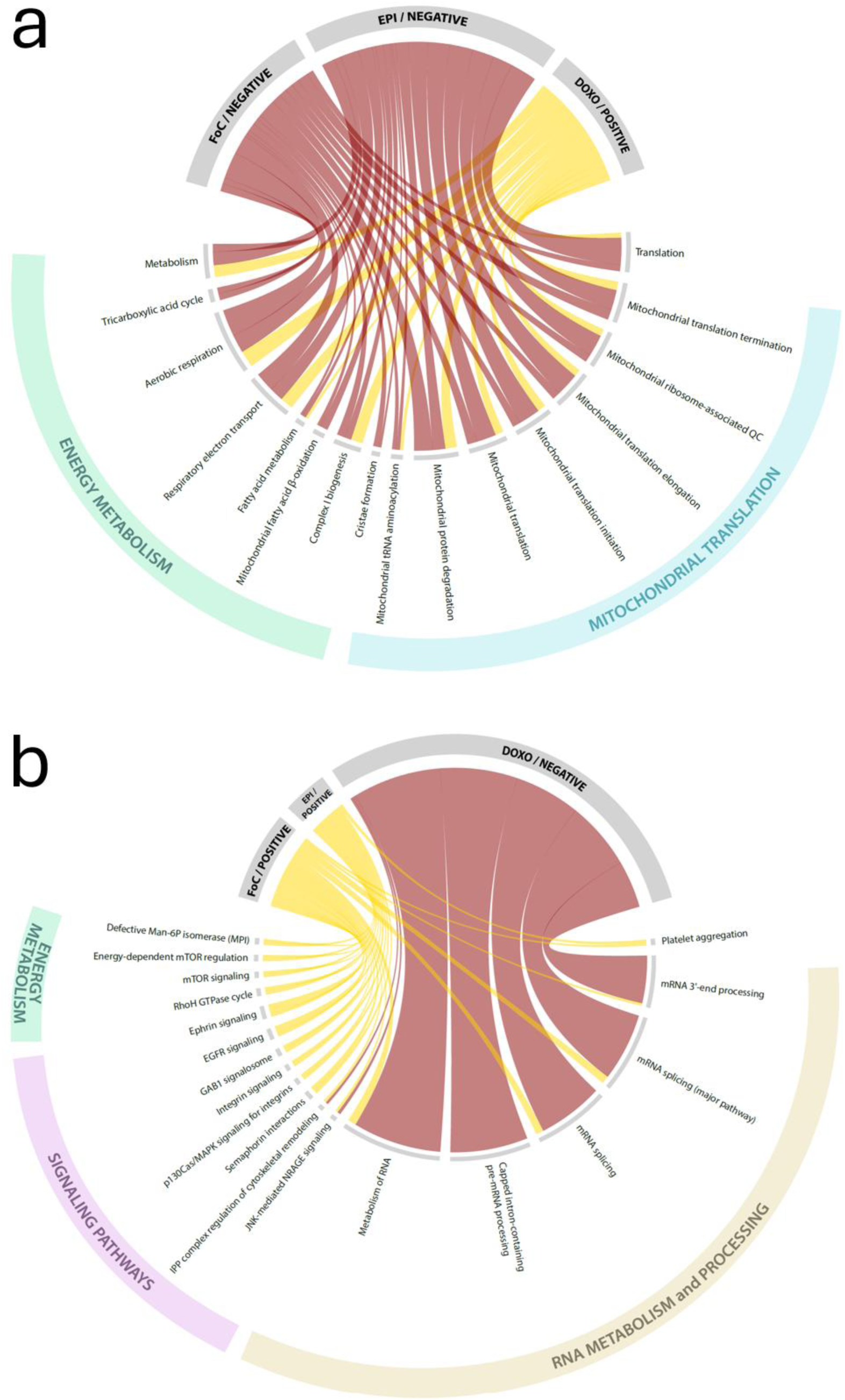
Chord diagrams summarizing significantly associated pathways shared across conditions. The chord diagrams depict the top biological pathways significantly associated with FoC (24 h post-treatment) and both drug treatments. (a) The first chord diagram depicts pathways with opposite directions of association: negative associations with FoC and EPI and positive associations with DOXO treatment (Pearson correlation). This reversal in correlation direction highlights the opposing system-level effects of the positive inotrope EPI and the negative inotrope DOXO on molecular processes associated with cardiac contractile function. (b) The second chord diagram depicts pathways shared between pairs of conditions among those positively associated with FoC and EPI and negatively associated with DOXO. Unlike panel (a), no pathways were shared across all three conditions. Abbreviations: DOXO, doxorubicin; EPI, epinephrine; FoC, force of contraction; QC, quality control; TCA cycle, tricarboxylic acid cycle.

Conversely, a more limited overlap was observed among pathways positively associated with FoC and EPI and negatively associated with DOXO (Figure 5b). Pathways related to energy metabolism regulation, particularly mTOR signaling pathway, were positively associated with both FoC and EPI but did not show a corresponding inverse association with DOXO. Similarly, pathways involved in cell signaling were positively associated with FoC and EPI, with only a limited subset (i.e. cytoskeletal remodeling regulation and JNK-mediated NRAGE signaling) showing negative associations with DOXO. Pathways related to RNA metabolism and processing were predominantly negatively associated with DOXO, with only mRNA splicing-related pathways positively associated with FoC. In addition to these pathways showing opposite or condition-specific associations, several pathways were associated with EPI and DOXO in the same direction despite their opposing inotropic effects. Among these, RNA metabolism and processing pathways showed negative associations with both EPI and DOXO, whereas several of the same pathways were positively associated with FoC. This overlap suggests that some molecular processes are commonly associated with drug exposure independently of the direction of the resulting contractile response.

Overall, the chord diagrams highlighted a prominent reversal in the association of energy metabolism and mitochondrial processes (e.g. mitochondrial translation and proteostasis) between FoC/EPI and DOXO. These processes therefore represent treatment-specific molecular responses associated with the divergent functional phenotypes induced by EPI and DOXO.

Although our TNE-based correlation approach does not determine pathway activity (activation or suppression, which would require further experimental investigation), it successfully captured differential molecular responses to drugs with opposite inotropic effects and identified specific cellular processes whose perturbations are directly associated with changes in cardiac function.

## DISCUSSION

In this study, we established and optimized a workflow combining sequential functional assessment and high-resolution proteomics in human engineered cardiac tissue models, enabling the quantification of over 7,000 proteins in each tissue subjected to functional characterization. This integrated workflow provides comprehensive molecular coverage while preserving the direct relationship between tissue function and proteome composition, allowing functional phenotypes to be interpreted in the context of their underlying molecular changes.

While tissue engineering and MS-based omics have rapidly advanced to replicate and characterize biological complexity, data analysis continues to rely largely on conventional statistical methods that analyze each data modality independently. This creates an increasing disconnect between experimental methods and analytical capability, particularly as modern biological studies increasingly generate heterogeneous datasets comprising functional measurements, molecular profiles and experimental metadata ^22,24,43^. Bridging these complementary layers of biological information therefore requires mathematical frameworks specifically designed to integrate complex, high-dimensional datasets rather than analyzing each modality in isolation ^40^. To address this challenge, we explored a mathematical framework based on TDA. Unlike conventional statistical approaches, TDA maps the underlying sample space using topological lenses derived from diverse data modalities ^27,29,36,37^. Rather than asking how heterogeneous datasets should be statistically combined, TDA shifts the analytical focus to the shape of the data space when viewed from complementary perspectives on the system state, and to its biological meaning ^29^. This enables direct cross-modal integration of fundamentally different data types while remaining robust to limitations frequently encountered in OoCs and engineered-tissue studies, including limited sample sizes, biological heterogeneity and high-dimensional datasets ^25^. The resulting unified mathematical model, the TCN, provides an intuitive and interactive representation of the integrated data that facilitates biological interpretation while preserving the complexity of the original measurements ^25^. Furthermore, because the TCN is generated independently of predefined biological assumptions and can integrate highly complex datasets, it supports the application of conventional statistical analyses. For example, our node-level enrichment approach enables correlation-based interrogation of molecular and functional relationships.

Our framework demonstrated how TDA can effectively integrate functional and proteomic information to resolve treatment-specific molecular responses in EHTs at the system-level. TNE-based correlations closely recapitulated correlations obtained from the complete dataset, supporting the robustness of the framework for preserving cross-modal relationships while organizing heterogeneous samples into biologically coherent subsets. Biologically, the pathway enrichment profiles associated with FoC closely mirrored those observed following EPI treatment, consistent with the positive inotropic effect of EPI. In contrast, DOXO displayed a markedly different enrichment profile, as expected for a drug with an opposing (negative) inotropic effect. These results highlight the ability of our framework to distinguish molecular responses associated with drugs exerting opposite effects on cardiac contractility. Through comparative analyses, energy homeostasis and mitochondrial processes emerged as the main cellular processes differentially associated with the divergent functional phenotypes induced by the two treatments. The chord-diagram analysis further highlighted these differences by revealing a pronounced reversal in pathway-phenotype associations between the positive and negative inotropic conditions. Altogether, these findings support the capacity of the proposed framework to move beyond individual protein-level associations and identify biological processes associated with distinct functional responses, although this does not establish whether the identified pathways are activated or suppressed.

An important consideration is the type of biological information provided by the data collected in this study. Expression proteomics provides a broad overview of protein abundance, however it does not directly capture protein activity or function. Changes in protein abundance therefore cannot be assumed to reflect proportional changes in enzymatic activity, signaling output, or pathway flux. Thus, expression proteomics should primarily be considered a hypothesis-generating approach for identifying candidate biological processes and pathways, which require validation through targeted functional assays. Pathway enrichment analysis has additional limitations, as databases such as Reactome are hierarchically organized and contain overlapping annotations, often mapping one protein to multiple pathways. Moreover, pathway nomenclature does not always accurately reflect the underlying biological process, particularly when proteins are shared across diverse physiological and disease-associated pathways. Throughout this study, these limitations were mitigated by interpreting enriched pathways as broader functional processes rather than only as independent entities. Nevertheless, pathway enrichment remains primarily a tool for pathway discovery and requires further functional validation.

Overall, this study demonstrated the ability of our TDA-based framework to integrate data from advanced human cardiac tissue models and high-resolution proteomics through topology-based analysis, enabling system-level characterization of drug responses and pathway discovery. Topology-based frameworks may therefore provide a scalable strategy for integrating functional and molecular data from increasingly complex engineered tissue models, supporting systems-level mechanistic studies and next-generation drug development.

## CONCLUSION

Understanding how molecular alterations give rise to functional phenotypes remains a major challenge when studying complex biological systems like human tissues. Here, we demonstrated how topological data analysis (TDA) provides a new mathematical framework for mathematical, cross-modal integration of proteome-wide alterations with functional readouts. Our framework enables molecular and phenotypic information to be interrogated within a unified system-level representation and, through our topological enrichment approach, to calculate correlation between numerical and categorical data. Besides enabling cross-modal data integration, TDA addresses limitations of standard statistical approaches when handling complex, heterogeneous, and high-dimensional data, while avoiding modality-specific preprocessing and distributional assumption. We also benchmarked our novel approach, topological node-level enrichment (TNE), which maps proteomics features, functional readouts, and categorical metadata onto the integrated network for quantitative cross-modal correlation. In human cardiac tissue as a test case, our framework identified cellular processes directly and oppositely associated with the divergent cardiac responses (altered contraction) to doxorubicin (negative inotrope) and epinephrine (positive inotrope), including energy homeostasis, mitochondrial translation and proteostasis. Beyond this application, our TDA-driven framework has been designed to be modular and scalable and is suitable for the integration of cross-modal proteomics and functional data in a range of *in vitro* tissue models, thus advancing systems-level studies for preclinical drug development.

## SUPPLEMENTARY MATERIALS & DATA AVAILABILITY

Supplementary figures, files and datasets have been deposited in the FigShare repository (FigShare.com), under the https://doi.org/10.6084/m9.figshare.33187935, and are publicly available as of the publication date.

The supplementary material includes:

- Supplementary Figure 1: Four dimensionality reduction methods applied to the entire sample-set (including 0h and 24h)
- Supplementary Figure 2: Heatmap of pairwise Pearson correlations between metadata using TCN node-level enrichment
- Supplementary Figure 3: Pairplot illustrating metadata relationships using TCN node enrichment
- Supplementary Figure 4: Customized weighted Voronoi graphs visualizing the results of the pathway enrichment analysis for the untreated samples
- Supplementary Figure 5: Customized weighted Voronoi graphs visualizing the results of the pathway enrichment analysis for the DMSO-treated samples
- Supplementary Table 1: Results of the node-level correlation analysis across conditions using different correlation thresholds
- Supplementary File 1: Settings and details of the proteomics data acquisition and annotation (Proteoscape)
- Supplementary File 2: Interactive topological connectivity network (TCN)
- Supplementary File 3: Sample composition of TCN nodes
- Supplementary File 4: Results of the pathway enrichment analysis (all conditions)
- Supplementary File 5: Full-scale Voronoi graph for FoC (negatively correlated features)
- Supplementary File 6: Full-scale Voronoi graph for FoC (positively correlated features)
- Supplementary File 7: Full-scale Voronoi graph for EPI condition (negatively correlated features)
- Supplementary File 8: Full-scale Voronoi graph for EPI condition (positively correlated features)
- Supplementary File 9: Full-scale Voronoi graph for DOXO condition (negatively correlated features)
- Supplementary File 10: Full-scale Voronoi graph for DOXO condition (positively correlated features)
- Supplementary File 11: Full-scale Voronoi graph for DMSO condition (negatively correlated features)
- Supplementary File 12 Full-scale Voronoi graph for DMSO condition (positively correlated features)
- Supplementary File 13: Full-scale Voronoi graph for UNTREATED condition (negatively correlated features)
- Supplementary File 14: Full-scale Voronoi graph for UNTREATED condition (positively correlated features)
- Supplementary Dataset 1: Force of contraction dataset
- Supplementary Dataset 2: Proteomics dataset
- Supplementary Dataset 3: Topological node enrichment (TNE) dataset
- Supplementary Dataset 4: Node-based correlations between proteins and metadata

## ACKNOWLEDGEMENTS

This study was financially supported by the Stofwisselkracht research grant 2023 (“HoPE” project to FC and HJCTW), Prinses Beatrix Spierfonds (“Op weg naar therapie” grant W.OR22-14 to FC), the European Research Council (“Heart2Beat” grant no. 101098372 to RP).

Figures 1a,b,c and 3a were generated using Biorender (www.biorender.com).

## ABBREVIATIONS

3D) EHT: (3-dimensional) engineered heart tissue
2-CAA: 2-chloroacetamide
ABC: ammonium bicarbonate
CFs: cardiac fibroblasts
CMs: cardiomyocytes
dia-PASEF: default long gradient data independent acquisition / parallel accumulation serial fragmentation
DMSO: dimethyl sulfoxide
DOXO: doxorubicin
DPBS: Dulbecco’s phosphate-buffered saline
DTT: dithiothreitol
EPI: epinephrine
FDR: false positive rate
FoC: force of contraction
hiPSCs: human induced pluripotent stem cells
LFQ: label-free quantification
MBR: match-between-run
MS: mass spectrometry
OoC: organ-on-chip
PCA: Principal component analysis
PCoA: principal coordinates analysis
PDMS: polydimethylsiloxane
PHATE: potential of heat-diffusion for affinity-based transition embedding
TCN: topological connectivity network
TDA: topological data analysis
timsTOF: trapped ion mobility spectrometry quadrupole time-of-flight mass spectrometer
TNE: topological node enrichment
UMAP: uniform manifold approximation and projection

